# Intermediate structural hierarchy in biological networks modulates the fractal dimension and force distribution of percolating clusters

**DOI:** 10.1101/2021.06.01.446578

**Authors:** Benjamin S. Hanson, Lorna Dougan

## Abstract

Globular protein hydrogels are an emerging class of materials with the potential for rational design, and a generalised understanding of how their network properties emerge from the structure and dynamics of the building block is a key challenge. Here we computationally investigate the effect of intermediate (polymeric) nanoscale structure on the formation of protein hydrogels. We show that changes in both the cross-link topology and flexibility of the polymeric building block lead to changes in the force transmission around the system, and provide insight into the dynamic network formation processes.

---

A vast range of hierarchical networks exist throughout biology, both natural and artificial, each with different macroscopic network properties emerging from the structure and dynamics of the subunits from which they are formed[1–3]. Naturally evolved examples include fibrin networks involved in blood-clotting[4], collagen networks contained in connective tissues and bones[5], and the actin and cytoskeletal networks which support cell structure and rearrangement[6, 7], among many others[8, 9]. Collagen has many types, but we find it useful to highlight that the hierarchical organisation of persistent collagen fibres[10, 11] forming mammal skeletons (together with calcium phosphate) gives them a high tensile strength at large length scales, and sufficient transverse flexibility to support the forces applied by musculature[12, 13]. Fibrin, on the other hand, forms a much more branched and isotropic network when part of a clot[14, 15] which, together with its native extensibility, enables it to elastically deform without fracture, and retain sufficient porosity to allow enzymes and antibodies to diffuse through[14, 16]. The scientific community have developed many artificial substitutes for biological networks. In addition to the polymeric networks we see in biology, which are artificially reproduced as semi-flexible polymer networks and gels[2], we also have peptide assemblies and gels[17, 18], colloidal gels[19, 20] and folded protein-based hydrogels[21–23]. Each have unique viscoelastic behaviours, finding applications in food science[24], as drug delivery systems[25, 26] and as candidates for artificial extra-cellular matrices and scaffolds[27–31]. However, it is not yet clear how sub-unit properties translate across the scales to generate the macroscopic responses we measure in practise. If we are to rationally design such systems, a cross-length scale understanding of hierarchical translation is necessary, and a key challenge in biological and soft matter physics[2].

In this letter, we investigate the use of short (5-mer) globular domain polymeric chains as a subunit in the simulated growth of biological networks. Viewed as an expansion of previous computational work related to pseudo-colloidal networks[32], we define these short polymeric subunits as “intermediate structure”, as they allow us to probe the differences in network formation dynamics between systems with spherical subunits (small aspect ratio), which may represent colloids or folded proteins, and polymeric systems (large aspect ratio). These 5-mers were designed such that at the volume fraction used in our simulations, the expected end-to-end distance of the polymers is approximately equal to the inter-particle distance at equilibrium[33] and thus, their local rotational (and translational) motion should become more restricted as their flexibility decreases due to the steric interference of neighbouring polymers. Short and finite-length polymeric assemblies have been investigated previously with respect to many of the referenced types of biological network[27, 34, 35], but here our focus is specifically aimed at the possibility of specifically designed folded polyprotein hydrogels. Polyproteins are used routinely in in single-molecule experiments[36–42] and have therefore previously been investigated as a viable design parameter for hydrogels[43]. It has also been suggested that polyprotein hydrogels may enable ensemble protein unfolding studies if correctly characterised[44]. We show that in the context of protein hydrogels these polymeric subunits provide novel, tunable single-molecular properties that propagate to the network level, enabling us to modulate the cross-link density, fractal dimension and force distribution throughout the network. At the same time, by comparison with related monomeric network formation simulations[32], they give theoretical insight into general dynamic principles of network formation.

Our simulations were performed with BioNet, a bespoke software package previously used to investigate the emergence of physical behaviour in both polymers[45] and protein-based hydrogels[32]. BioNet models proteins as coarse-grained spheres with dynamics calculated via a Brownian dynamics protocol. The objects are able to sterically interact using an efficient volumetric potential, and can be connected with Hookean springs. Importantly, cross-linking sites are explicitly defined at the surface of each object, with each sphere having local rotational degrees of freedom explicitly modelled. As such, connections between objects combined with the steric potential in BioNet generate a realistic emergent stiffness[45]. When two unbound cross-link sites come within 0.3nm of one another in our simulations, a stiff, Hookean cross-link dynamically forms. When combined with the steric potential, then, a space-filling network emerges. As we are representing protein hydrogels, spheres with explicit cross-linking sites represent proteins with specifically engineered tyrosine residues that enable photo-chemical cross-linking[46, 47]. Further details on the method can be found in our previous work[32, 45].

The parameter space of subunit structure and flexibility is shown in Fig. 1a). With insight from our previous work[45], we parametrised these globular domain polymers with three different stiffnesses as measured by the persistence length, *L_p_*, all with approximately the same contour length *L_c_*. The three stiffnesses are defined by *L_p_/L_c_* ≈ 0.5, *L_p_/L_c_* ≈ 1.0 and *L_p_/L_c_* ≈ 1.5 for the “Rigid”, “Semi-flexible” and “Flexible” 5-mers respectively. Each sphere along the polymer has radius *R* = 2.5nm, and spheres along the polymer were connected by permanent Hookean springs with equilibrium length *l* = 0nm. 5-mers were chosen to match previous experimental[43] and theoretical[44] work on characterising their potential as polyprotein hydrogels. *R* = 2.5nm was chosen to represent bovine serum albumin (BSA), a protein previously simulated using BioNet[48]. The cross-link site topology was varied by progressively increasing the number of sites along the polyprotein. As shown in Fig. 1a), the polymer model with the smallest number of sites (M_0_) has 4 sites (1 on node P_0_, 2 on node P_2_, and 1 on P_4_), the minimum number for a 3D topological subunit (a tetrahedron)[49]. These sites span the entire length of the polymer, so that while the cross-link site topology is constant, the *geometry* is affected by the flexibility of the polymer itself. As the total number of sites, *N_s_*, increases, moving from M_0_ to M_5_, we effectively fill up the remaining space along the polymer, starting at the centre and moving outwards. The cross-link topology can therefore be uniquely referred to by the value *N_s_*, but we keep in mind that the flexibility of the polymer can still affect how these sites are geometrically arranged in space. Prior to each network formation simulation, the simulation was populated with pre-assembled polymers and thoroughly equilibrated. All simulations contained *N* = 1000 polymers in a simulation box with periodic boundary conditions and side length *L* = 187nm, giving a volume fraction *f_v_* = 0.05. This volume fraction has previously been shown to form percolating networks for monomers[32] and hence should also form them with polymers due to their larger aspect ratio, especially given experimental observations of such systems[43]. As previously stated, *f_v_* = 0.05 also has the interesting property of making each polymer be at the very limit of immediate steric interaction with neighbouring polymers, so our systems can be viewed as being around the central point between colloidal and polymeric network behaviour. The box size was chosen to be significantly larger than the cluster size measured in the experimental work on polyprotein gels[43] and monomeric gels[46, 47] but interestingly, this makes our box of the same order as clusters recently characterised in locally reinforced monomeric BSA gels[48]. All analysis was performed after 30*μ*s of simulation time, and examples of the range of possible resultant network structure are given in Fig. 1b). Further methodological details are given in the Supplementary Information[33].

**FIG. 1:**
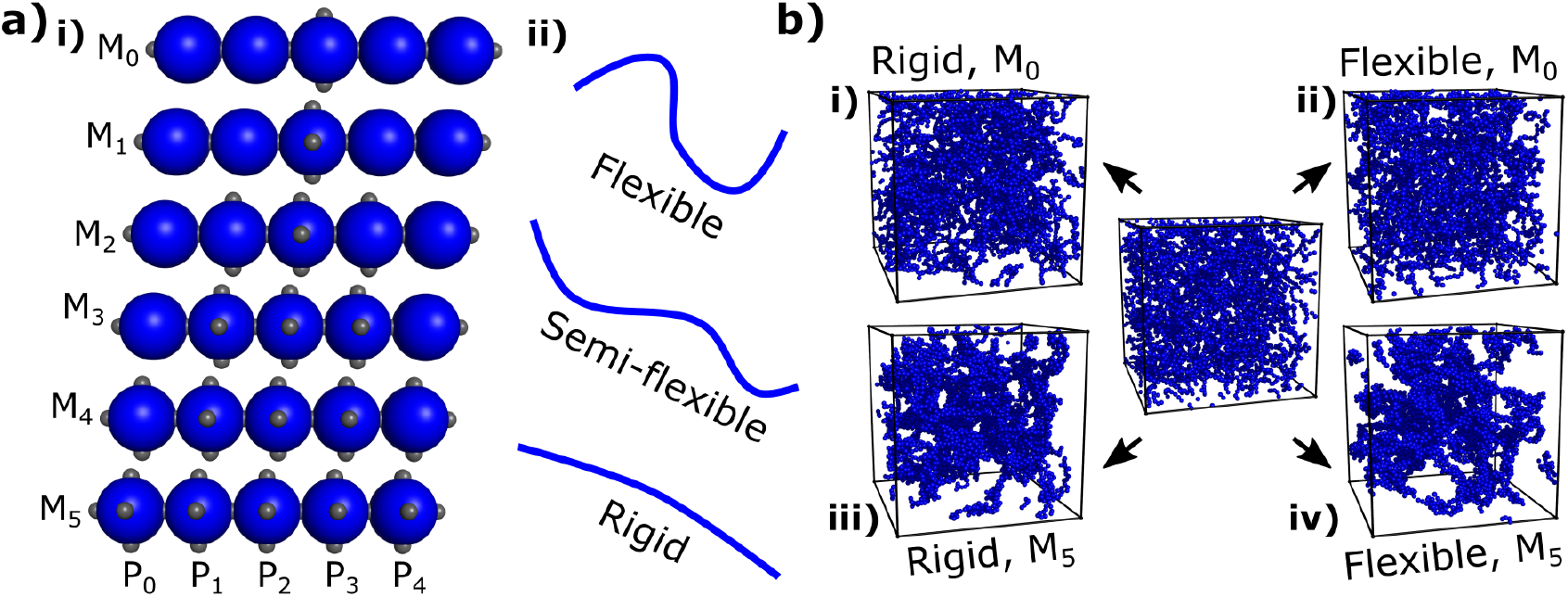
**a)** The parameter space explored by our simulations. **ai)** We alter the cross-link topology through models M0 to M5 by progressively increasing the number of cross-link sites along the globular polymer, as shown by the grey dots on each blue sphere labelled from P0 to P5. **aii)** We alter the polymeric flexibility by slightly changing the length of the linker domain connecting the spheres[45]. Each parameter can be independently altered, yielding 18 unique systems. **b)** Example networks showing the limiting cases of observed network structures. An example initial state (post-equilibration) is shown in the center, and final snapshots of limiting-case simulations are shown. **bi)** The most rigid polymer with model M_0_. **bii)** The most flexible polymer with model M_0_. **biii)** The most rigid polymer with model M_5_. **biv)** The most flexible polymer with model M_5_.

We begin with an investigation of the cross-link coordination per monomer and the network fractal dimension. These structural properties have been analysed in the context of many biological networks[50, 51], and identified as useful in the characterisation of polymeric networks[52]. We previously calculated these values in monomeric pseudo-colloidal networks[32] where both were shown to affect the pore size distribution and mechanical response of those networks. Fig. 2 shows the expected coordination per monomer, *C*, within the network, ranging from *C* = 0.52 ± 0.01 for the most rigid polymer with model M0, to *C* = 2.82 ± 0.01 for the flexible polymer with model M5. For the higher *N_s_* models we observe more cross-linking, just as in our previously characterised monomeric systems[32]. Even though cross-linking never reaches full saturation, it is still the case that cross-linking is restricted by the availability of sites. We also see that the more flexible the polymer, the higher the propensity for cross-linking. This indicates that there is an enthalpic cost associated with forming more cross-links in these polymeric networks. A direct comparison can be made with our previously studied monomeric networks to show the effect of including intermediate structure on the cross-linking. Consider our previous monomeric networks with *N_s_* = 6 per monomer. Pre-assembling these *N_s_* = 6 monomers into 5-mers results in model M5. For such monomers, we observed *C* = 3.55 ± 0.01[32], a value higher than even the most flexible of our polymeric systems. The additional enthalpic cost of this pre-assembly therefore leads to a lower cross-linking propensity, and the higher the enthalpic cost (the higher the rigidity), the lower the cross-linking. We can the reduce cross-linking even further by explicitly removing sites along the polymer. The more flexible polymers will still be able to deform to make the most cross-linking sites, but by the time we reach extremely limited cross-link topologies (models M0 and M1), there are simply no sites available. Subunit engineering, then, can be utilised to reduce cross-linking via two distinct mechanisms, via explicit removal of cross-linking sites, as is also the case with monomeric networks, or by introducing enthalpic energy penalties in the form of intermediate structure and mechanical response.

**FIG. 2:**
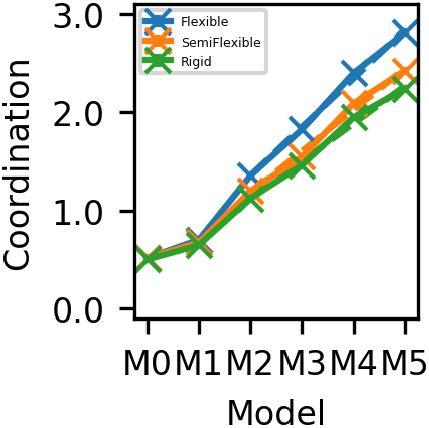
Expected coordination per monomer, not including the initial connections present due to polymeric pre-assembly. Flexible polymer systems shown in blue, semi-flexible in orange and rigid in green.

An important insight to the polymeric subunits is understanding specifically where these cross-links are formed along the polymeric contour. Fig. 3 shows the total expected cross-link coordination of each node (monomer) P0-P4 along the polymer chain, and also within each model, for the flexible and rigid subunit networks. These expectation values were calculated by averaging across each “equivalent” node within the network, and again only includes covalent cross-links, not the softer connections defining the polymer contour itself. The semi-flexible case is provided as Supplementary Information[33].

**FIG. 3:**
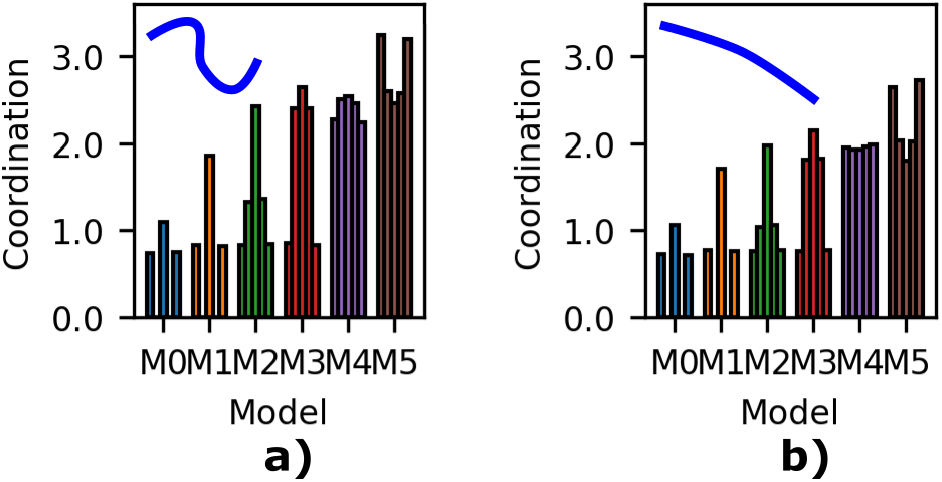
Expected coordination of each specific node P0-P4 along the polymer chain, not including the initial connections present due to polymeric pre-assembly. Within each model (separated by colour), each bar represents node P0-P4 from left to right. **a)** Flexible polymer systems **b)** Rigid polymer systems.

We again see that more sites to a monomer leads to more cross-links forming on that specific monomer, and that the more flexible polymers have more cross-links on each monomer, showing that the previous insights can be localised to each individual monomer within the polymer. Interestingly, as we move through our models from M0 to M5, increasing the number of sites on the inner monomers and moving outwards, the new cross-links do not displace those present on the outer nodes. In other words, in the flexible and rigid systems, the coordination on the outer nodes P0 and P4 changes only very slightly. However, as we move from model M2 to M3, adding additional sites to P1 and P3, the coordination on P2 changes as well. This indicates that the outer node cross-linking propensity is in some sense independent from the rest of the polymer. Normalisation of the local node coordination is shown in Fig. 4, where each coordination value is divided by the total number of available cross-link sites. Here we see that, per site, it is the outer-most nodes which are more likely to be cross-linked into the network. Insights from the following analysis of the fractal dimension analysis will elucidate a possible cause for this behaviour.

**FIG. 4:**
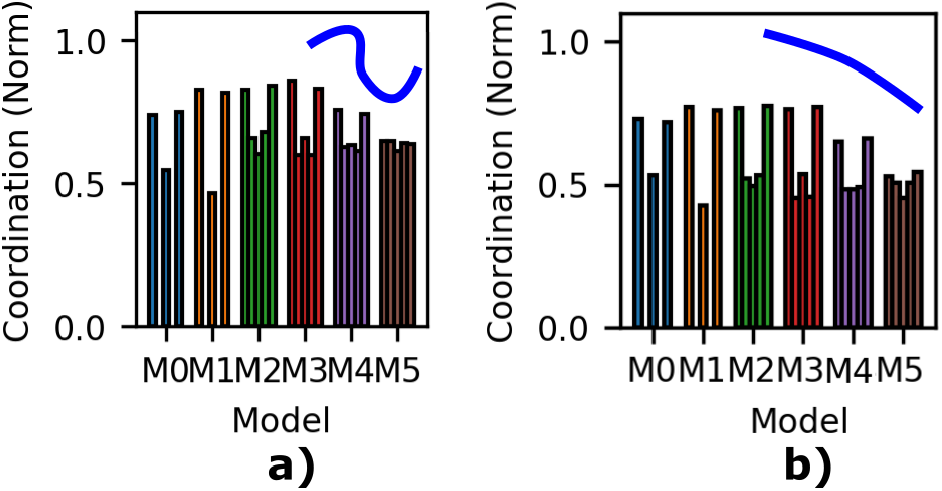
Normalised expected coordination of each specific node P0-P4 along the polymer chain, not including the initial connections present due to polymeric pre-assembly. Within each model (separated by colour), each bar represents node P0-P4 from left to right. **a)** Flexible polymer systems **b)** Rigid polymer systems.

Fig. 5 shows both the change in fractal dimension, *D_f_* and upper fractal limit, 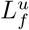, of the entire system as a function of polymeric flexibility and model, calculated via the box-counting technique detailed in the Supplementary Information[33]. Briefly, for a finite-sized system such as ours, it is not appropriate to think of the fractal dimension in terms of the infinitely recursive self-similarity it is usually associated with. Rather, the fractal dimension can be thought of here as a measure of how a finite amount of material is arranged in a finite volume i.e. the morphology of the system. A high *D_f_* indicates denser clusters, and a low *D_f_* indicates sparser clusters (i.e. more “branch-like”). Thus, from the perspective of a finite system, the fractal dimension is a valid measure only when specifically quoted between two limiting length-scales. In our case, the lower length limit, 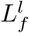 = 2.5nm, the protein radius, and the upper length limit, 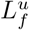, is measured as part of our box counting process. Outside of these limits, the system appears three-dimensional either due to a lack of resolution (above 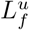) or a lack of further structure (below 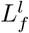). The upper fractal limit can therefore be thought of as a measure of structural homogeneity throughout the entire network. A high 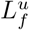 indicates inhomogeneity, as clustering must be sufficient to generate voids in the system larger than the inter-particle spacing at equilibrium. The converse is true for low 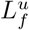, and the difference between Fig. 1bi) and Fig. 1bii), with respect to Fig. 5b) shows this clearly. Thus, for a finite system, we may expect that higher *D_f_* values (i.e. denser clusters) would be paired with higher 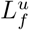 values, and indeed, this is exactly what we observe in Fig. 5.

**FIG. 5:**
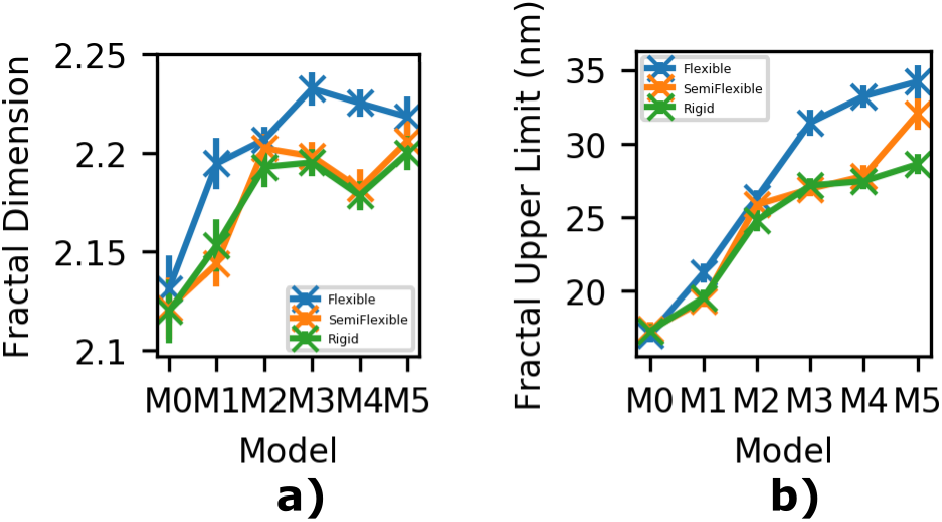
Fractal characteristics of our simulated networks. Flexible polymer systems shown in blue, semi-flexible in orange and rigid in green. a) Fractal dimension. b) Upper fractal limit.

With these insights, we interpret Fig. 5 as showing that more flexible polymers lead to denser clusters forming, as do higher *N_s_* models, with *D_f_* = 2.12 ± 0.02 and 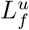 = 17.3 ± 0.5nm for the most rigid polymer with model M0, and *D_f_* = 2.22 ± 0.01 and 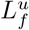 = 34.2 ± 1.1nm for the most flexible polymer with model M_5_. In the Supplementary Information, we show that these fractal characteristics both grow over the course of the simulation and additionally, that higher *N_s_* models have a significantly lower percolation lag time, indicating rapid percolation[33]. With respect to the previous cross-link characterisation, this suggests that as more sites become available, the flexibility of the polymers enable these sites to cross-link via cluster collapse, rather than cluster growth. In other words, because our polymers are long enough to interact simply via rotation, and because the cross-linking propensity of the outer-nodes P0 and P4 seem to be unaffected by flexibility, we infer that network percolation happens quickly in these polymeric networks, followed by network (cluster) collapse. In contrast, our previous monomeric networks, having a much smaller aspect ratio, likely form smaller clusters which locally collapse, and these clusters connect and grow into a percolating network. This is supported by the fact that while the range of fractal dimension values in our monomeric systems was approximately the same as in the polymeric networks, adding more sites to the monomers actually *reduced* the fractal dimension rather than increasing it[32], indicating that more sites in the monomer led to a more branched structure. Roberts *et al.*[47] make a clear distinction between glassy dynamic growth defining colloidal gels[53] and the nucleation, growth and connection of smaller clusters defining chemical gels[54]. We are suggesting that increasing the aspect ratio moves the network formation dynamics towards th glassy behaviour of colloids without increasing the volume fraction. Subunit engineering, then, can be used to alter not only the cross-linking propensity and fractal dimension characteristics (and thus the network inhomogeneity and porosity), but also the fundamental dynamic processes of network formation.

We now consider how these structural properties relate to the distribution of force around the networks. We calculate forces in each of the three Cartesian directions local to each node as shown in Fig. 6. These directions are defined by the initial node orientation, which rotates together with the node itself throughout the simulation. The “axial” direction is tangent to the local polymer contour, and each of the radial directions represent the directions perpendicular to the polymer contour.

**FIG. 6:**
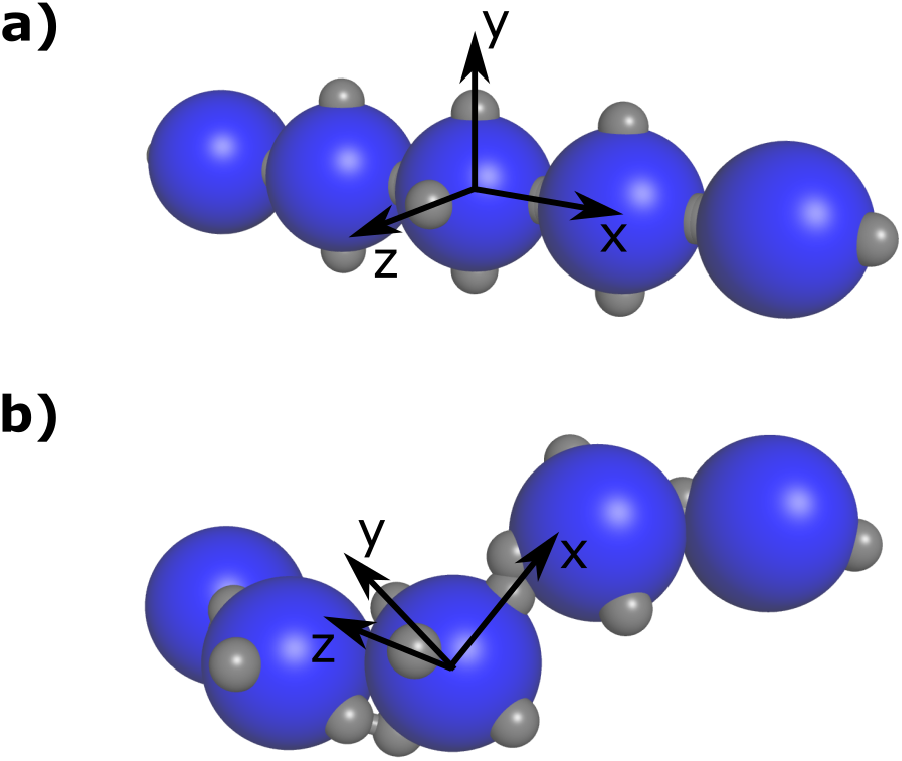
Definition of a local coordinate system for a node within a polymer. The local *x* axis is the “axial” direction, and *y* and *z* axes are the two “radial” directions. **a)** The initial coordinate system before any equilibration has taken place coincides with the global Cartesian coordinates. As such, cross-linking sites are not required to keep track of rotation. **b)** Some time through a BioNet simulation. As the node rotates, we keep track of the local coordinate system. It is clearly different for each node.

Fig. 7 shows the expected forces on each node along the polymer chain, in each of the directions for each model in the flexible and rigid systems. It is important to note that the forces on our nodes here are likely over-estimates, as the spheres are rigid bodies and unable to internally relax leaving the cross-links themselves as the sole enthalpic components. However, this limitation of physical realism allows us to observe the immediate internal forces in the network by analysing the cross-links themselves as they apply force to the spheres, and where force would be applied under swelling or external stress, without having to model more complex sphere deformations and stress. As such, a qualitative interpretation of the following section is appropriate.

**FIG. 7:**
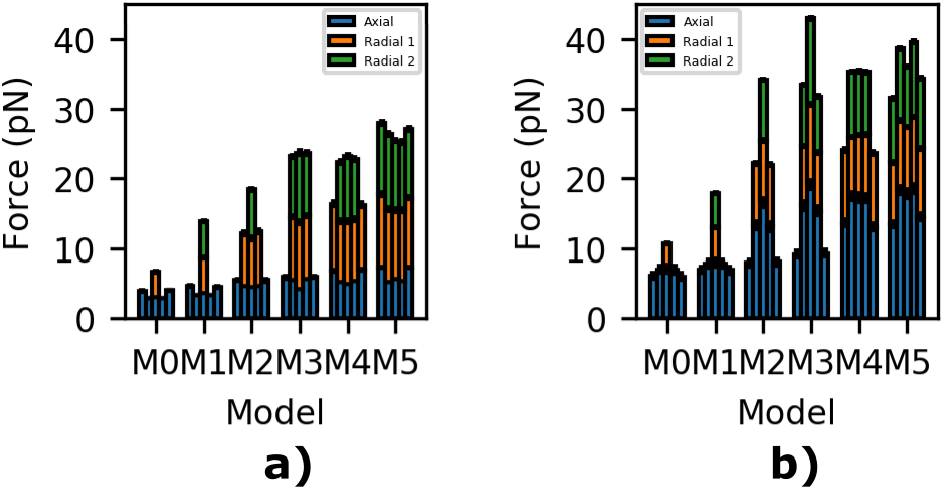
The expected forces on each specific node P0-P4 along the polymer chain in the three Cartesian directions local to the polymer. The “axial” direction, shown in blue, is along the polymer contour, and the two “radial” directions, shown in orange and green, are the mutually orthogonal directions defined by the cross-linking sites (if present). **a)** Flexible polymer systems. **b)** Rigid polymer systems.

We see that the total force on the rigid objects (the sum of all forces) are each greater than their flexible equivalent, and generally as the number of cross-link sites increases (and hence, the coordination), so does the total force. In the flexible systems the forces are generally larger in the radial directions, which shows the difference between the cross-links and softer interactions along the polymer contour. In a system with deformable spheres, we would expect the strain would therefore occur primarily between cross-links, as there is much more mechanical flexibility and conformational freedom along the polymeric contours. In other words, assembling soft, flexible intermediate structures enables stress relaxation along the polymeric contours. However, as the polymer becomes more rigid the converse is true. It is very interesting to note the much higher force on the central node P2 of with model M3 compared with M5 in the rigid model. This implies that the specific network configuration of this model prevents stress relaxation throughout the network, although it is not entirely clear why.

With respect to the increase in *N_s_* with the models, we see that in both the flexible and rigid systems, it is not just the radial forces that increase with *N_s_*, even though the axial bonding remains constant throughout all simulation. The addition of cross-links in the radial direction increases the force in the axial direction too, dramatically in the case of the rigid systems. This shows that the connectivity itself, the limitation of conformational freedom, affects not only how much force is applied to each monomer, but how force is distributed along each polymer and by extension throughout the network. However, there is still a coupling with the with the coordination, as we see that the force on the central nodes decreases with respect to the outer nodes as *N_s_* increases. Thus, intermediate substructure enables us to tune not only the cross-linking behaviour but also the stress relaxation pathways throughout the network. Summarising this results presented, we envisage the pre-assembly of monomeric subunits into either polymers, or perhaps a more exotic form of intermediate structure, to be a unique, useful mechanism for both experimental tuning of network structure & mechanical response, and also a new coordinate for exploring the dynamics of network formation.

## Supporting information

Extended supplementary information on methods, results and discussion

## ACKNOWLEDGMENTS

We thank Kalila Cook for useful discussions on the fractal dimension of discrete, rigid networks. We acknowledge the EPSRC funding through grant EP/P02288X/1 awarded to Professor Lorna Dougan.

