## Extended supplementary information on methods, results and discussion for "Intermediate structural hierarchy in biological networks modulates the fractal dimension and force distribution of percolating clusters"

#### DATA ACCESS & REPRODUCIBILITY

All of our polyprotein network simulations were performed using BioNet, a software package in development at the University of Leeds. This software is available for download from the Bitbucket repository `https://bitbucket.org/GokuBH/proteinhydrogelsoftware/`. Although the core software is in active development for additional applications and features, for reproducability purposes the branch *PolyproteinNetworkSims* remains unchanged since this work was performed.

All data and graphs used in this work, as well as additional movies, can be found at NOTE: DOI.

#### PERSISTENCE LENGTH PARAMETRISATION

To investigate the effect of persistence length on network growth, we first require BioNet polymers which have an appropriate range of persistence lengths. For a given contour length $L_c$, we require three polymers with persistence lengths $L_p$ such that $L_p/L_c \sim 0.5$ ("Flexible"), $L_p/L_c \sim 1.0$ ("Semi-flexible") and $L_p/L_c \sim 1.5$ ("Rigid"). Using the results of our previous investigation into globular domain polymers[45] we performed a series of simulations of 10-mers, where each spherical domain had radius $R = 2.5$nm and the linker domains between had an equilibrium length $l = 0.0$nm. To vary the value of $L_p$, we changed the stiffness of the linker domain via the spring constant $k$. We previously showed that increasing this parameter reduces the thermal length fluctuations $\Delta l = \sqrt{k_B T/k}$, and thus increases the local steric interactions between spherical domains, leading to an overall increase in $L_p$. Biophysically, increasing $k$ approximately corresponds to an reduction in the number of amino acids making up the linker domain, a locally flexible worm-like chain polymer which therefore has approximately harmonic end-to-end fluctuations.

From these simulations (which ran for hundreds of Rouse times for statistical convergence), Fig. S1a shows the set of explicitly calculated correlation traces using the equation:

$$\langle \vec{t}(s) \cdot \vec{t}(s') \rangle = \exp\left(\frac{|s' - s|}{L_p}\right), \quad (1)$$

where $s$ and $s'$ are distances along the polymer contour, and $\vec{t}$ is the normalised gradient vector at that point. We see that each curve shows the characteristic exponential decay as expected, with a reduced persistence length as the normalised value $\Delta l/2R$ increases. Extracting these $L_p$ values and plotting them against $\Delta l/2R$ gives us Fig. S1b, to which we have applied an empirical exponential fit. From here, we used the fit to calculate the value of $\Delta l/2R$ required for a given $L_p/L_c$, and the values we require are shown as dashed lines extended to each axis.

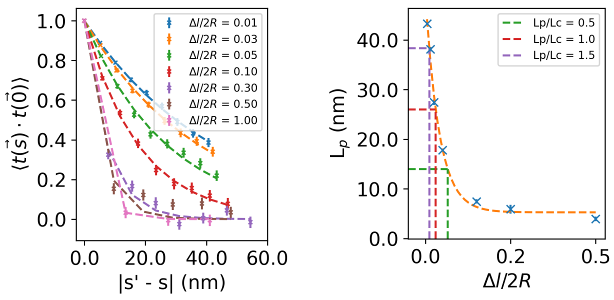


(a) (b)

FIG. S1: Data from the parameter sweep of linker stiffness values, leading to variations in the polymer persistence length. **a)** Gradient vector correlation traces. **b)** Extracted persistence lengths as a function of normalised linker fluctuations, a stand-in for the linker stiffness.

| $\Delta l/2R$ | $k$(pN/nm) | $L_c$(nm) | $L_p$(nm) | $\mathbf{L_p/L_c}$ |
|---|---|---|---|---|
| 0.132 | 9.46 | 27.64 | 15.01 | **0.54** |
| 0.061 | 44.13 | 26.22 | 25.22 | **0.96** |
| 0.023 | 305.62 | 25.46 | 38.62 | **1.52** |

TABLE S1: The final persistence length data calculated from models representative of what will be used in the main body of work.

We used each of these $\Delta l/2R$ values to parametrise three further simulations to verify our approach, and reanalysed them using the same approach to calculate the persistence length. These final simulations 5-mers, as they are what will be used in the main body of work. The results are given in Table S1.

As $\Delta l$ is a fluctuating coordinate (due to the application of thermal noise), the effective $L_c$ calculated from each simulation is slightly different, leading to a slight deviation in our $L_p/L_c$ values. However, they are still reasonably close to our initial aims and so we can justifiably parametrise our network polymers using these values of $k$ to give distinguishable flexible, semi-flexible and rigid polymers.

#### ROTATIONAL RESTRICTION OF STRUCTURAL SUBUNITS

With our previous monomeric networks, a monomer was able to freely rotate about its own centroid until a cross-link formed, at which time local monomeric rotation was restricted. As clusters get larger and larger, eventually the cluster will be unable to rotate without coming into contact with other objects with which to interact. With polymeric systems, however, the initial polymer is so long that rotational motion is restricted immediately. Indeed, a major insight of the tube model in a polymer melt is that reptation is a dominant form of dynamic motion[55], almost the conceptual opposite of pure rotation. So where do our short polymers fit into this? For a given volume fraction $f_v$, a structural subunit has a certain amount of available volume, $v_a = v_s/f_v$, where $v_s$ is the volume of the subunit. Our subunit is a globular domain polymer, and in isolation this polymer would freely rotate in 3D space and trace out a volume. This volume can be approximately characterised by a sphere with a radius $R_{rms} = \frac{1}{2}\sqrt{\langle E^2 \rangle}$, where $\langle E^2 \rangle$ is the expected square of the end-to-end distance. Using the standard expression for the end-to-end distance[55], it can be shown that for our polymers, $R_{rms} = A\sqrt{2L_cL_p}$, where $A = \frac{1}{2}\sqrt{1 - \frac{Lp}{Lc}\left(\exp\left(-\frac{L_c}{L_p}\right)\right)}$ is some prefactor dependent on the stiffness of the polymer.

We can now say that if $R_{rms} < \frac{1}{2}v_a^{\frac{1}{3}}$, then the subunit is initially free to rotate, otherwise it is hindered by the presence of other subunits in the system. Expanding all these terms and making the relative stiffness of our polymer, $L_p/L_c$, the subject, we obtain an inequality defining whether the polymer is free to rotate

$$\frac{L_p}{L_c} < B\left(\frac{1}{f_v N_m^2}\right)^{\frac{2}{3}} \tag{2}$$

where $N_m$ is the number of monomers in a polymer, and $B = 0.598$ for our flexible polymers, $B = 0.859$ for our semi-flexible polymers and $B = 1.213$ for our rigid polymers. Substituting in the relevant values from the main text into Eq. 2, $f_v = 0.05$ and $N_m = 5$, we would expect that each of our systems are in fact rotationally restricted at the volume fraction used in the main work (Eq. 2 is not satisfied), albeit only slightly. In the case of the flexible polymer case, especially, both sides of Eq. 2 are almost equal, and in each case are very close to the exact point of transition between networks in which the initial subunits have rotational freedom, and networks where they do not. This is likely a key characteristic of the transition from globular, colloidal networks to long polymeric networks, and may explain why our semi-flexible and rigid structural characterisations are so similar in comparison to the flexible system.

### CALCULATING THE FRACTAL DIMENSION BY BOX COUNTING

To calculate the fractal dimension of our networks, we employ box counting. By sub-dividing our simulation box into tessellating boxes, with $N_x$ boxes along each edge, then we can count the total number of boxes $N$ which contain any protein material. For small $N_x$, we would expect every box to be filled as we are at low spatial resolution. However, as we increase $N_x$, there will be some boxes which fit into the gaps in the network and hence we would expect fewer boxes to be filled. In the limiting case of $N_x \to \infty$, $N$ and $N_x$ are related through the volume fraction, but our analysis does not go this far. We scan our simulation frames and count $N$ for each sub-division $N_x$ and fit the following piece-wise function to the data:

$$\log_2(N) = \begin{cases} 3\log_2(N_x) & N_x \leq N_x^t \\ D_f\left(\log_2(N_x) - \log_2(N_x^t)\right) + 3\log_2(N_x^t) & N_x > N_x^t \end{cases} \tag{3}$$

where $N_x^t$ is the transition point between regimes. Fig. S2 shows the data for both the flexible and rigid (polyprotein) sets of data.

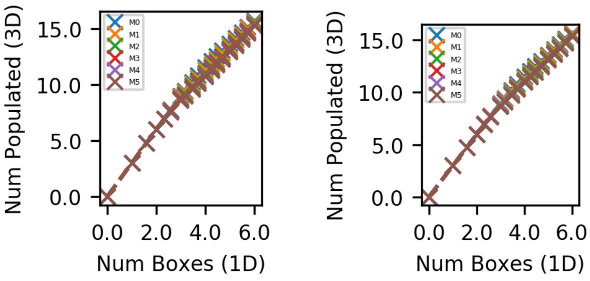


(a) (b)

FIG. S2: Data comprising the box counting method for calculating fractal dimensions. **a)** Flexible polyproteins. **b)** Rigid polyproteins.

### SEMI-FLEXIBLE POLYMER NETWORKS

Fig. S3 shows the local coordination properties of the semi-flexible polymer systems, and Fig. S4 their localised force distributions.

### DYNAMIC EVOLUTION OF STRUCTURAL CHARACTERISTICS

Fig. S6 shows the evolution over time of maximum cluster size for each system.

To each of the traces in Fig. S6 we fit a logistic curve to extract dynamic time parameters. Fig. S7 show the central logistic value $t_0$ and the growth rate *tau* for each polymeric flexibility. The parameter $t_0$ is related to the percolation lag time, and thus we can see that the presence of more sites both significantly reduces the lag time

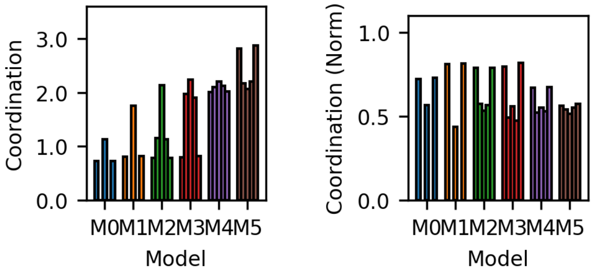


(a) (b)

FIG. S3: The coordination properties of the semi-flexible polymer systems. **a)** Total coordination per node **b)** Normalised coordination per node.

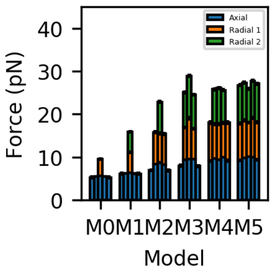


FIG. S4: The localised force distribution of the semi-flexible networks.

and also increases the growth rate once percolation has been achieved.

Figs. S8, S9 and S10 show the fractal dimension $D_f$ and upper fractal limit $L_f^u$ for each of the polymeric flexibilities.

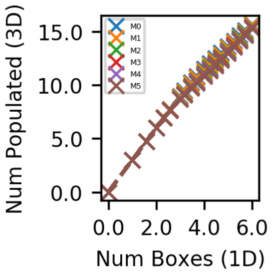


FIG. S5: Data comprising the box counting method for calculating fractal dimensions in the semi-flexible polymer networks.

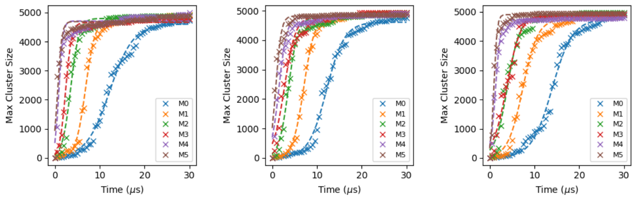


(a) (b) (c)

FIG. S6: Dynamic evolution of the maximum cluster size over time. **a)** Flexible polymer networks. **b)** Semi-flexible polymer networks. **c)** Rigid polymer networks.

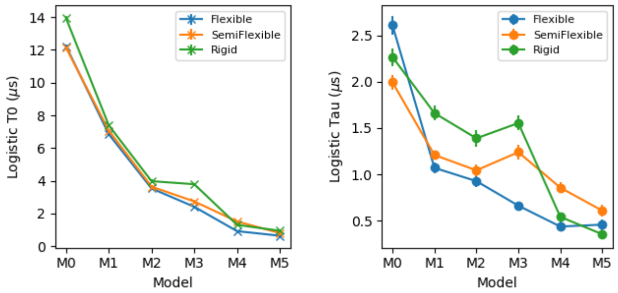


(a) (b)

FIG. S7: Logistic curve fitting parameters for the cluster growth traces of the polymer networks. **a)** Central logistic time $t_0$. **b)** Logistic growth rate $tau$.

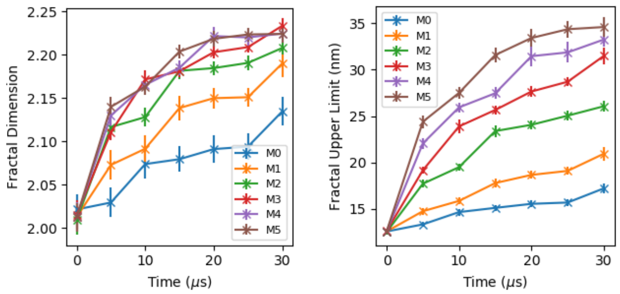


(a) (b)

FIG. S8: The fractal characteristics of the flexible polymer networks. **a)** Fractal dimension. **b)** Fractal upper limit.

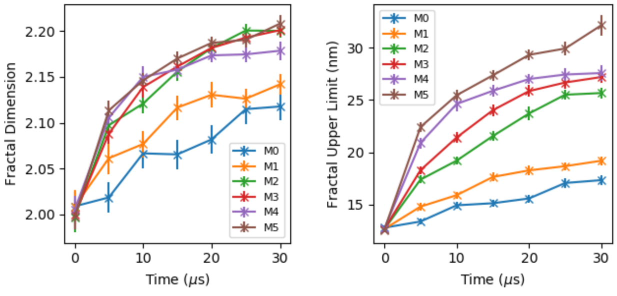


(a) (b)

FIG. S9: The fractal characteristics of the semi-flexible polymer networks. **a)** Fractal dimension. **b)** Fractal upper limit.

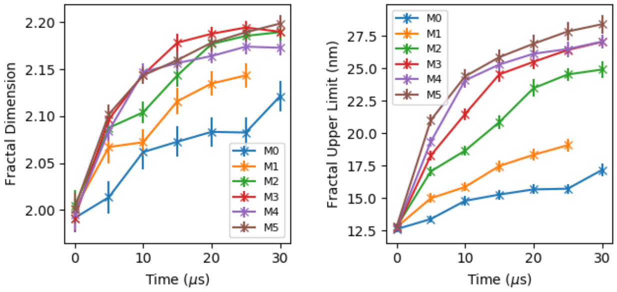


(a) (b)

FIG. S10: The fractal characteristics of the rigid polymer networks. **a)** Fractal dimension. **b)** Fractal upper limit.
